# Error propagation in constraint-based modeling of Chinese hamster ovary cells

**DOI:** 10.1101/2020.07.09.195594

**Authors:** Diana Széliová, Dmytro Iurashev, David E Ruckerbauer, Gunda Koellensperger, Nicole Borth, Michael Melcher, Jürgen Zanghellini

## Abstract

**Background:** Chinese hamster ovary (CHO) cells are the most popular mammalian cell factories for the production of glycosylated biopharmaceuticals. To further increase titer and productivity and ensure product quality, rational systems-level engineering strategies based on constraint-based metabolic modeling, such as flux balance analysis (FBA), have gained strong interest. However, the quality of FBA predictions depends on the accuracy of the experimental input data, especially on the exchange rates of extracellular metabolites. Yet it is not standard practice to devote sufficient attention to the accurate determination of these rates.

**Results:** In this work we investigated to what degree the sampling frequency during a batch culture and the measurement errors of metabolite concentrations influence the accuracy of the calculated exchange rates and further, how this error then propagates into FBA predictions of growth rates. We determined that accurate measurements of essential amino acids with low uptake rates are crucial for the accuracy of FBA predictions, followed by a sufficient number of analysed time points.

**Conclusions:** We observed that the measured difference in growth rates of two cell lines can only be reliably predicted when both high measurement accuracy and sampling frequency are ensured.

## 1 | INTRODUCTION

Chinese hamster ovary (CHO) cells are the primary host for the production of biopharmaceuticals, particularly monoclonal antibodies and other complex therapeutic proteins^1^. Their main advantages include the ability to perform human-like post-translational modifications, in particular glycosylation, suspension growth in serum-free chemically defined media and a low risk of viral infections, which makes them a safe host for the production of human therapeutic proteins^2,3^. However, the development of new producer cell lines is a time-consuming, costly and laborious process, based mostly on laboratory evolution and high-throughput screening of thousands of clones. It often takes several months to obtain a high producer and this trial-and-error process must be repeated for each new product^4^. Furthermore, it is not clear which factors limit product formation^5^.

As a result of the success of metabolic modeling in designing microbial cell factories^6^, interest has grown in applying these methods for the analysis and optimisation of CHO as well^7^. However, applications of modeling to CHO cells remain limited as key (bioinformatic) resources became available only recently. The publication of CHO’s genome sequence^8^ enabled the *in silico* reconstruction of its metabolism^9^. Along with recent updates^10,11^ these genome-scale metabolic models sit at the heart of constraint-based modeling approaches that allow to computationally connect genotype and phenotype^12^. In fact, such a genome-scale metabolic model coupled with a model of the secretory pathway^13^ validated gene knock-outs that led to increased productivity and growth^14^. Hence, genome-scale modeling has great potential to improve CHO cell line development by identifying bottlenecks in productivity and designing engineering strategies to resolve these.

One of the most common metabolic modeling approaches is constraint-based modeling, such as flux balance analysis (FBA)^15,16^, especially for analysis of genome-scale metabolic models^17,18^. The accuracy of FBA predictions depends on the correct reconstruction of the metabolic pathways, the biomass composition and the uptake and secretion rates of extracellular metabolites which are used as constraints^16,19^. Previously we showed that the quality of growth rate predictions by FBA in CHO depends mostly on the accuracy of the measured uptake and secretion rates^20^. While simple organisms such as bacteria or yeast can grow on a single carbon source (thereby simplifying the analysis), mammalian cells grow in complex media containing numerous essential metabolites and secrete several byproducts. Typically, more than 20 exchange rates have to be measured in order to perform FBA, which makes the analysis much more challenging than for simpler organisms on minimal media. However, despite the importance and the difficulty of determining accurate exchange rates for mammalian cells, little attention has been paid to this topic so far.

To determine exchange rates accurately, it is necessary to measure the metabolite concentration throughout the cultivation at a sufficient sampling frequency. In the literature, the concentrations of extracellular metabolites are commonly measured once per day, typically resulting in 4-6 time points in total^21,22,23,24,25,26,27,28^. However, this might not be enough to obtain accurate uptake and secretion rates.

Here we investigated the impact of sampling frequency and the error of metabolite concentration measurements on the calculation of exchange rates and subsequently on FBA predictions of growth rate. We determined which exchange rates have the biggest impact on FBA and what accuracy and sampling frequency is required to detect a given difference between two cell lines.

## 2 | METHODS

The simulations of experimental data, statistical analysis and visualisation were done in R version 3.4.3^29^. FBA^15,16^ was performed in python 3.7.4 using package COBRApy^30^ with Gurobi solver. All data and scripts were deposited in Mendeley Data at https://data.mendeley.com/datasets/5vn5m33wpr/draft?a=3e7afba4-42e0-45b4-9e16-7adb419251f9.

### 2.1 | Reference FBA

To generate a reference state, FBA was performed with the experimental data and genome-scale metabolic models taken from Széliová et al.^20^ for eleven out of thirteen available datasets (two datasets (`GScd4-8mMCD` and `DXepo-0mMCD`) were omitted from the analysis due to very inaccurate growth rate predictions). The experimentally measured specific uptake and secretion rates of glucose, amino acids, lactate and ammonium were used as constraints (see Table A1 for two example cell lines – `K1par-8mMAP` and `HYher-8mMCD`; the full dataset can be found in Mendeley Data, see link above). Cysteine and tryptophan uptakes were left unconstrained due to the lack of quantitative experimental data. Biomass production was set as the objective function (reaction `“R_biomass_specific”`). Then, growth rate was fixed to the predicted value and the uptakes of cysteine and tryptophan were minimized to obtain estimates for their uptake rates. Lactate secretion was left unconstrained for the cell line `DGpar-8mMCD` as in the original paper (otherwise the predicted cysteine uptake rate was unrealistically high – higher than the glucose uptake rate). The predicted growth rates were used as reference states in the subsequent analyses.

### 2.2 | Simulation of concentration profiles

The experimental exchange rates from Széliová et al. ^20^ (see Table A1 and Data Mendeley), the predicted growth rates and the uptake rates of cysteine and tryptophan from the reference FBA (see Section 2.1) were used to simulate concentration profiles using Equation (1),

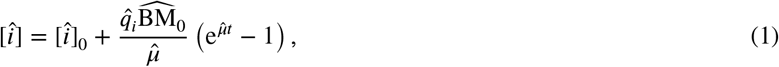

where 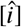 is an ideal concentration of metabolite *i* during exponential phase, 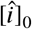 is the initial concentration of metabolite *i* in the medium, 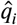 is the reference exchange rate, 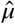 is the growth rate, *t* is time of cultivation and 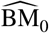 is the initial amount of biomass, calculated from an initial cell concentration (1.6 × 10^5^ viable cells/mL) and the experimentally measured dry mass per cell (`K1par-8mMAP`: 252.3 pg/cell, `HYher-8mMCD`: 279 pg/cell, see Data Mendeley for the full dataset). The values for 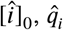, cell dry mass and initial cell concentration were taken from Széliová et al.^20^ (see Table A1 & Mendeley Data). 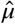 is the growth rate predicted by the reference FBA. The time of cultivation corresponds to the length of exponential phase, which was set to 90 hours (the mean length of exponential phases of the two cell lines). Samples were generated at regular intervals of 6, 12 or 24 hours or – to resemble typical working shifts – at irregular intervals with 4 samples per day – every 4 h with a 12 h gap or every 2.5 h with a 16.5 h gap (the gap was positioned either in the beginning of each day or after the first four “dense” sampling time points, e.g. hours 0, 12, 16, 20, 24, 36 etc. or 0, 4, 8, 12, 24, 30 etc.).

To simulate experimental data, normally distributed noise was added to the ideal concentrations

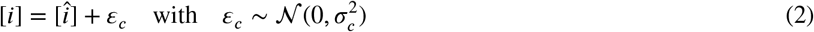

where *ε_c_* is a standard normal random variable with relative standard deviation (RSD) 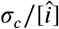 chosen between 0.02 and 0.20. Note that the noise increases with the concentration level. The range of the RSDs was chosen based on the available data from Széliová et al.^20^, where median RSDs of the concentration measurements from 3-6 replicates were 1.7-7.1% (mean RSDs 5.3-19.8%). 1000 concentration profiles were generated for each metabolite. Afterwards, an equation of form (1) was fitted to each profile with R function `nls`, using *q_i_* and [*i*]_0_ as the fitting parameters, thereby obtaining 1000 sets of exchange rates. Standard errors (SEs) of the fitted rates were used as a measure of accuracy. For a graphical representation of the workflow see Figure A1, Sim 1.

In the second version of the simulations, biomass concentrations were perturbed, in addition to concentration profiles. For this, cell concentration data BM was simulated with Equation (3),

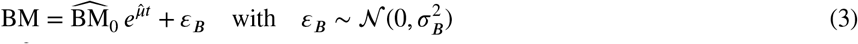

where *t* is time of cultivation (0-90 h), 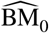 is the initial biomass (the same as in Equation (1)) and 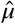 is the growth rate from the reference FBA. Normally distributed noise of 2-20% 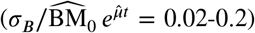 was added in the same way as described above for the concentration profiles. The parameters *μ* and BM_0_ were estimated by fitting the simulated biomass concentrations with an exponential growth function using R function `nls`. Note that the estimated growth rate (*μ*) was used for the simulations of metabolite concentrations with Equation (1) (instead of the growth rate predicted by the reference FBA) (Figure A1, Sim 2).

### 2.3 | FBAs with simulated exchange rates

The simulated exchange rates (*q_i_*) and the SEs were used as constraints for FBA. The lower and upper bounds of the exchange reactions were set to *q_i_* – SE_*i*_ and *q_i_* + SE_*i*_, respectively. In rare cases, it was not possible to fit Equation (1) to the concentration profiles due to high noise. In those cases, the exchange rates were left unconstrained. Biomass production was used as the objective function (reaction `“R_biomass_specific”`).

## 3 | RESULTS

First, we defined a reference state for further simulations. We used experimental data and genome-scale metabolic models of eleven CHO cell lines/conditions^20^ in the exponential phase of a batch culture. FBA was performed based on experimental data as described in Methods. The uptake reactions of cysteine and tryptophan were computationally estimated due to the lack of quantitative experimental data. We maximized the biomass production and obtained growth rates 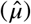 in the range 0.0263-0.046 h^−1^. These were considered as reference or “true” values for the subsequent simulations. Except for DXB11 models, the predicted uptake rates of cysteine and tryptophan exactly corresponded to the requirements for the synthesis of biomass and were used for further simulations. (In the DXB11 models cysteine is not essential and the predicted cysteine uptakes were lower than the biomass requirement.)

### 3.1 | Low exchange rates are highly inaccurate

First, 1000 sets of concentration profiles of 23 extracellular metabolites (amino acids (AAs), ammonium, glucose, and lactate) were simulated by adding normally distributed, independent random errors with a mean of zero and a RSD ranging from 2% to 20% to all reference distributions. Three replicates were simulated (representing three independent experiments, e.g. three bioreactor runs) and the distributions were sampled at three regular intervals - every 6 h, 12 h or 24 h (corresponding to 16, 8 or 4 time points throughout the batch, respectively). Figure 1 exemplarily shows simulated histidine concentration profiles for sampling intervals of 6 h, 12 h, and 24 h and RSDs of 2%, 10%, and 20%.

**FIGURE 1.**
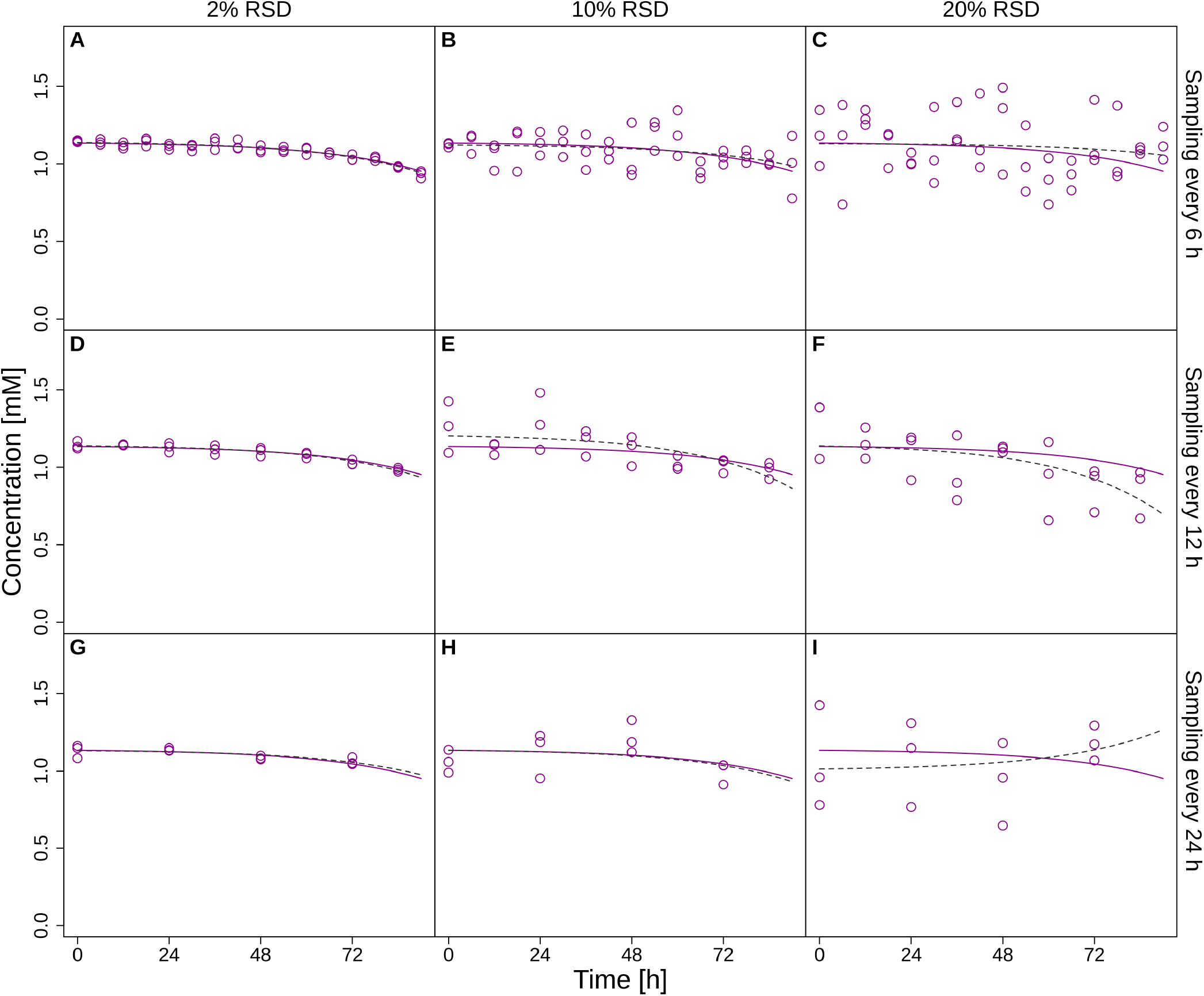
Simulated reference profiles (solid lines) and sampled perturbed concentrations (points) of histidine during the exponential phase of a batch cultivation at a sampling frequency of 6 h (panels A, B, C), 12 h (panels D, E, F) or 24 h (panels G, H, I) with 2% (panels A, D, G), 10% (panels B, E, H) or 20% (panels C, F, I) RSD of the replicate concentration measurements. Dashed lines represent fits to the data using Equation (1). Data is shown for cell line `K1par-8mMAP`.

Next, we calculated exchange rates by fitting Equation (1) to the simulated concentration profiles and analytically evaluating its derivative. Low sampling frequency and high error can result in bad fits that in some cases may even predict secretion instead of consumption (e.g. see Figure 1, panel I).

We used the SEs to assess the accuracy of the calculated exchange rates (*q_i_*). Even though the simulated concentration data has the same RSD for all metabolite concentrations, the relative standard errors (RSEs) of the calculated exchange rates (SEs divided by the absolute value of the exchange rates, *q_i_*) markedly vary. More specifically, the closer the exchange rate is to zero, the higher the RSE and the wider the distribution (Figure 2, Figure A2). Figure 2 a shows examples for a sampling frequency of 6 h (purple colours), where the simulated RSD of the concentrations is 2% and the medians of the RSEs of the rates are 1% for glutamine (high uptake), 2% for asparagine (medium uptake) and 6% for histidine (low uptake). 10% RSD of concentration measurements leads to median RSEs of the rates of 5%, 10% and 29% for glutamine, asparagine and histidine, respectively. If the sampling interval is increased from 6 h to 24 h (green colours), the resulting RSEs of the exchange rates triplicate. For instance, with a 10% concentration error, the median error in the uptake of histidine increases up to 86% and the distribution has a very long tail. Figure A3 shows the relationship between RSDs of the metabolite concentrations and RSEs of the calculated exchange rates at different sampling frequencies for cell line `K1par-8mMCD`.

**FIGURE 2.**
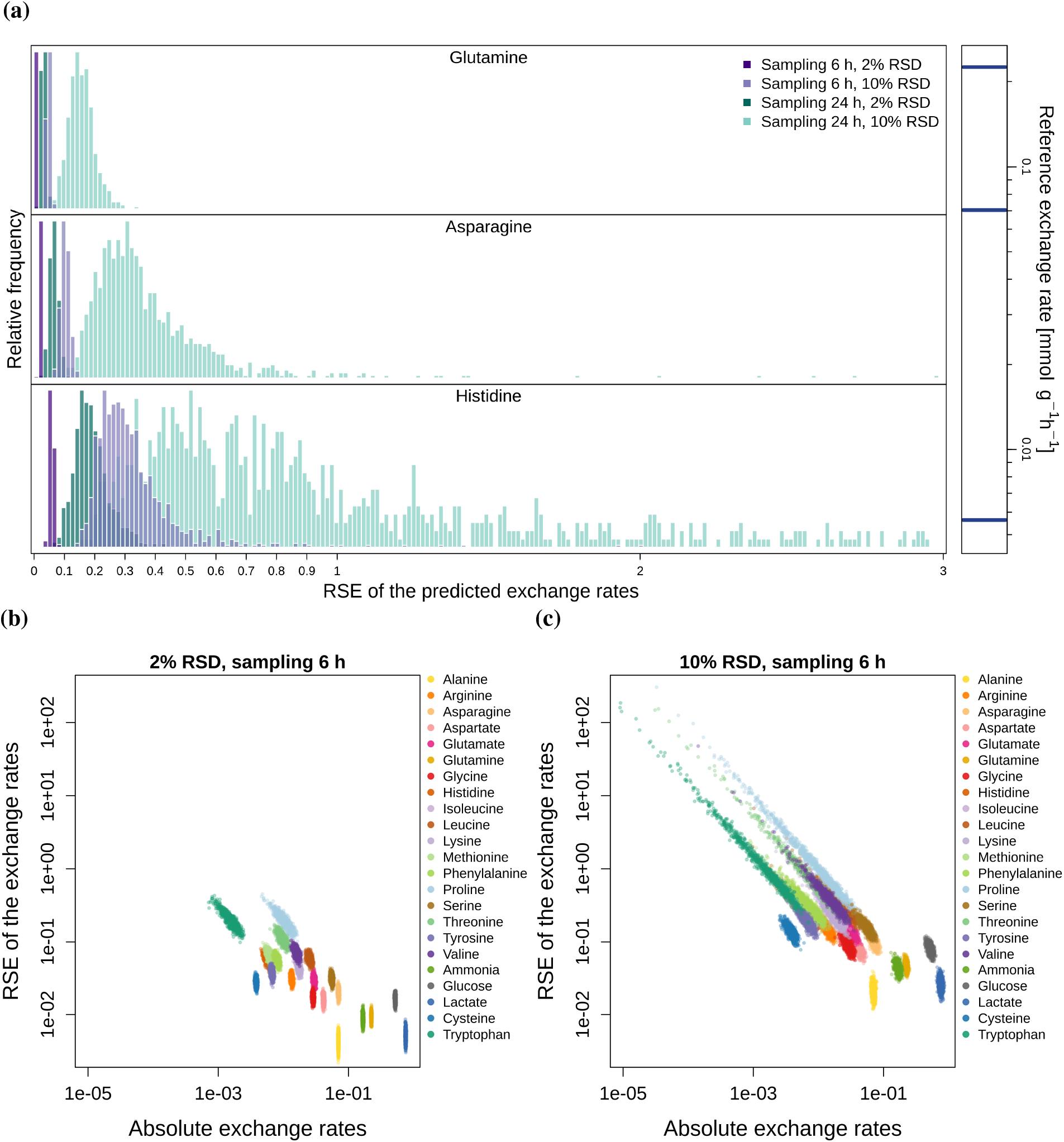
Panel a: distribution of the RSEs of three example uptake rates (for cell line `K1par-8mMAP`). The bars on the right side indicate the magnitudes of the uptake rates (top, glutamine; middle, asparagine; bottom, histidine). The lower the exchange rate, the higher the RSE and the wider the distribution. Purple colours show the distributions for sampling frequency of every 6 h, green colours for frequency of every 24 h, each for 2% and 10% RSD of the concentration measurements. Bottom: RSEs of the exchange rates as a function of the absolute values of exchange rates shown for 2% (panel b) and 10% (panel c) RSD of the concentration data.

### 3.2 | Growth predictions are strongly sensitive even to small metabolite concentration errors

We constrained the genome-scale metabolic model iCHO1766 with the computed exchange rates [*q_i_ -SE_i_, q_i_*+SE_*i*_] and predicted maximal growth rate with FBA. Figure 3 shows the ratios between predicted and reference growth rates for two selected cell lines in different media (`K1par-8mMAP` and `HYher-8mMCD`). As expected, the distribution of growth rates gets wider with the increasing RSD in the metabolite concentrations and decreasing sampling frequency. At RSD of 20% and sampling frequency of 24 h, the predicted growth rate varies between zero and three times the true value. Note that the distribution gets skewed because the growth rates cannot be negative. Figure A4 displays the RSDs of the predicted growth rates as a function of concentration RSDs, showing that the data follows a square root law. That means that even small concentration errors lead to significant deviations of the growth rate predicted by FBA. However, growth rate RSD for concentration RSD of 20% is expected to be only twice higher than the one for concentration RSD of 5%.

**FIGURE 3.**
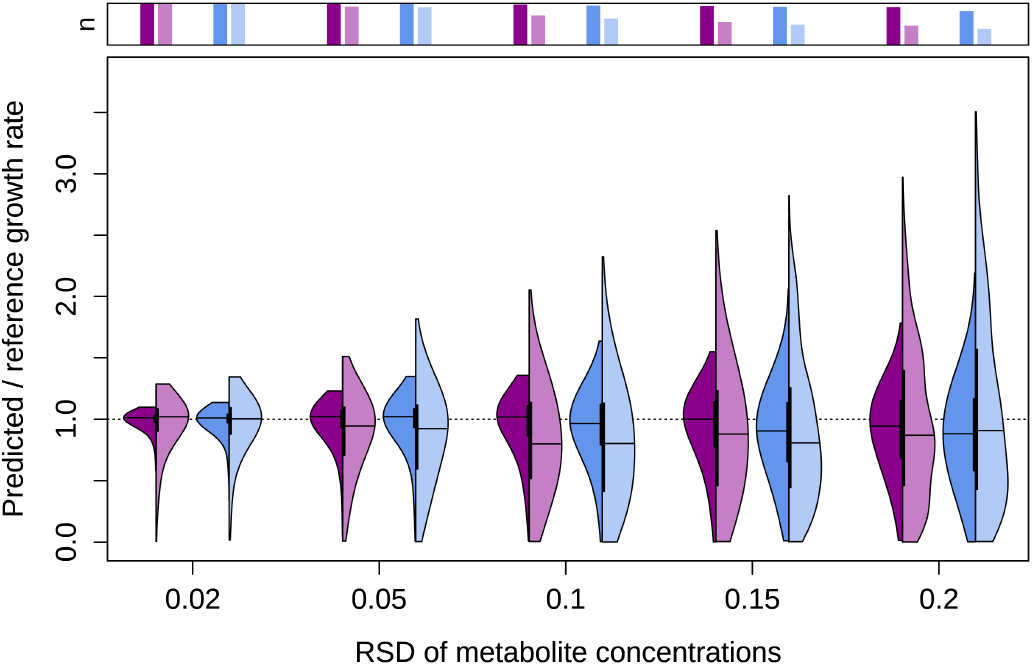
FBA predictions at different concentration RSDs for two cell lines (purple: `K1par-8mMAP`, blue: `HYher-8mMCD`). The left side of the violin plots corresponds to sampling every 6 h, the right side to every 24 h. The barplots above the plots indicate the fraction of feasible FBA solutions. The apparent cutoffs on the top at lower RSDs are artifacts due to the visualization.

Furthermore, with increasing RSD and decreasing sampling frequency we observe that there is a growing number of infeasible FBA problems (indicated by the bars on top in Figure 3). At RSD of 20% and sampling frequency of 24 h, more than half of the associated linear problems are infeasible. The predictions for the slower cell line (`HYher-8mMCD`, blue colours) are consistently worse, which can be explained by the on average lower uptake rates, which increases the RSEs of the calculated uptake rates (Figure A2).

Apart from the metabolite concentrations, the calculation of exchange rates takes the growth rate *μ* as an input (see Equation (1)). To check the impact of the error of the cell concentration measurements on the calculations of the exchange rates and the FBA predictions, we added noise not only to the metabolite concentrations but also to the cell concentrations, where we introduced 6% RSD (the accuracy of Vi-CELL XR (Beckman Coulter)^31^, an automated cell counting device commonly used to quantify cell concentration). Figure A5 d shows that the predictions are practically indistinguishable from the results in Figure 3. Growth rates are typically high enough (0.02-0.04 h^−1^)^20^ and thus can be determined with high accuracy. To verify this, we analysed the variation in the estimated growth rates at 2-20% measurement RSDs (see Figure A5 a and A5 b). Even if the measurement error is 20% and sampling only once per day, the RSDs calculated from 1000 estimated growth rates were below 15%. At the selected measurement RSD of 6%, the RSDs of the estimates were only 2-3%. Furthermore, we compared the RSDs of the estimated exchange rates with or without perturbing the cell concentration by 6%. Figure A5 c shows the maximum observed differences between exchange rate RSDs estimated with or without perturbing the cell concentration. Based on these results, we concluded that measurement errors of the growth rate have only a small effect on our simulations and need not be considered further.

Another variable in Equation (1) is the cell dry mass (contained in BM_0_). However, this parameter is commonly determined only once for each particular cell line and reused for all further experiments, so the error in this parameter was not considered and the value was fixed to the previously measured cell line/condition spedific values^20^.

### 3.3 | Essential amino acids determine the growth rate predictions by FBA

Next, we determined which exchange rates have the largest influence on growth rate predictions with FBA. Based on Figure A3, we hypothesized that the metabolites with largest relative error have the largest impact. Thus, if those were measured accurately, it would improve the prediction accuracy. To test this, we constrained the RSD of the concentrations to 2% for the top seven metabolites with highest RSEs of the rates. This strongly improves the prediction accuracy as measured by a decrease in the interquartile ranges for `K1par-8mMAP` (before/after): 0.13/0.09, 0.25/0.12, 0.37/0.16 and 0.47/0.2 for 5-20% RSDs, respectively (see Figure 4 a). Note that for 2% RSD, the results are the same, because in both cases all metabolites have 2% RSD. The seven most error-prone uptake rates were uptakes for proline, tryptophan, threonine, valine, methionine, leucine and histidine, the last 6 of which are essential components. The results for the cell line `HYher-8mMCD` show a similar trend (see Figure A6).

**FIGURE 4.**
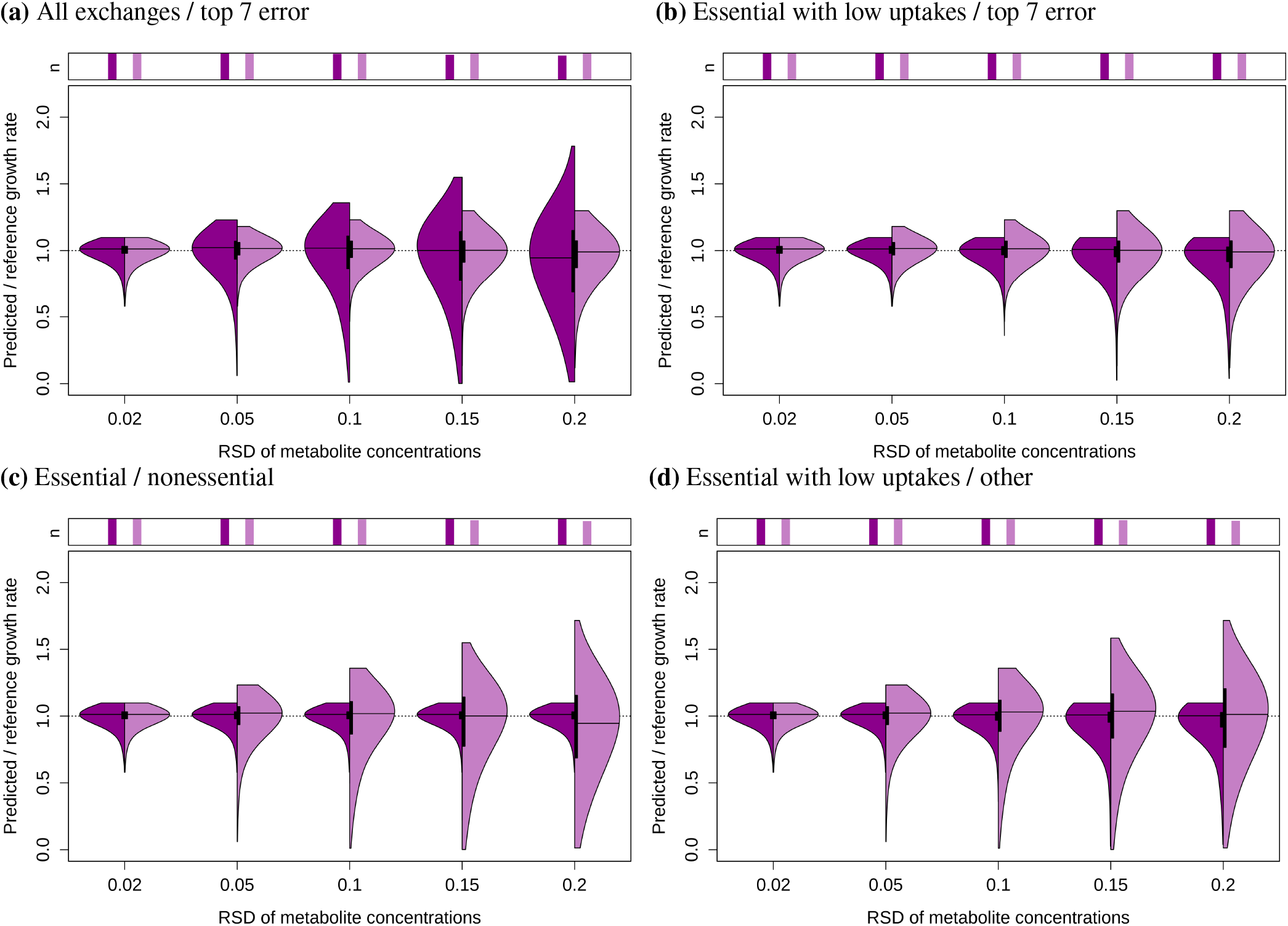
FBA predictions when different groups of metabolites are measured accurately. In both panel a and b, the right halves of the violin plots show results when the RSDs of metabolites with top 7 highest RSEs of the rates are set to 2%. In panel a, the left halves show results when RSDs of all metabolites are varied (the same as the left halves in Figure 3) and in panel b when RSDs of the AAs with low normalized uptake rates are set to 2%. In panel c, in left halves, the RSDs of the essential AA concentrations are set to 2%, while all others are varied (group 1 constant/group 2 varied); in the right halves, the other way around (group 1 varied/group 2 constant). In panel d, in left halves, the RSDs of essential AAs with uptakes < 1.5× the biomass requirements are set to 2%, while all others are varied (group 1 constant/group 2 varied, the same as the left halves in panel b); in the right halves, the other way around (group 1 varied/group 2 constant). For simplicity, the data is shown only for cell line `K1par-8mMAP` at the sampling frequency of every 6 h. The barplots above the plots indicate the fraction of feasible FBA solutions. The apparent cutoffs on the top are artifacts due to the visualization.

As the uptake of essential AAs cannot be compensated by the uptake of other AAs, we next tested whether growth rate is limited mainly by the uptake rates of essential AAs. Therefore, we split the exchange rates into two groups - group 1: uptake rates of essential AAs (histidine, isoleucine, leucine, lysine, methionine, phenylalanine, threonine, tryptophan, valine, cysteine, arginine) and group 2: all other uptake and secretion rates. We varied the RSD of the concentration data for one group, while keeping the RSD of the second group constant. When the error of the essential AA concentrations is kept constant (group 1), varying the error of the nonessential AAs has no effect on the FBA predictions (Figure 4 c and Figure A6 c, left halves). On the other hand, increasing the error of the essential AA concentrations leads to a large increase in the variability of the FBA solutions (Figure 4 c and Figure A6 c, right halves) and also has a minor effect on the number of infeasible FBA problems.

The uptake rates and biological functions of the essential AAs vary. Some of them are used only for the generation of cell biomass (see next paragraph), while others are also metabolized for other purposes (e.g. generation of energy), so we expected that their impact on the FBA predictions might differ. Therefore, the next grouping was based on the biomass requirements of the essential AAs, which can be calculated as growth rate × AA coefficient in the biomass equation. Again, the exchange rates were divided into two groups: group 1: essential AAs with uptake rates smaller than 1.5× their biomass requirement (for cell line `K1par-8mMAP`: arginine, lysine, phenylalanine, threonine, tryptophan, cysteine; for cell line `HYher-8mMCD`: arginine, leucine, lysine, methionine, phenylalanine, tryptophan, cysteine); group 2: all other exchange rates. Figure 4 d and Figure A6 d show that varying the concentration errors of AAs in group 1 (right halves), which are used only for biomass formation, has a much bigger impact on FBA predictions than varying the errors of metabolites in group 2 (left halves). Conversely, it makes only a tiny difference whether or not we keep the error of group 2 at 2% or leave the exchanges unconstrained (right halves of Figures 4 d and A6 d vs. Figure A7). Finally, no such effects were apparent when we repeated the same procedure with random groupings (see Figure A8).

Together this data demonstrates that the quality of FBA predicted growth rates is (primarily) determined by the uptake rates of essential AAs, which are used mostly for generation of biomass. The set of the relevant AAs is cell line-specific (according to the interquartile ranges, Figure 5 a). However, measuring them accurately leads to a larger improvement than accurate measurements of the high-error AAs (Figure 4 b). Note that only two out of six AAs for `K1par-8mMCD` (threonine, tryptophan) and three out of seven for `HYher-8mMCD` (leucine, methionine, tryptophan) from the “low uptake” group are also in the “top 7 high errors” group.

**FIGURE 5.**
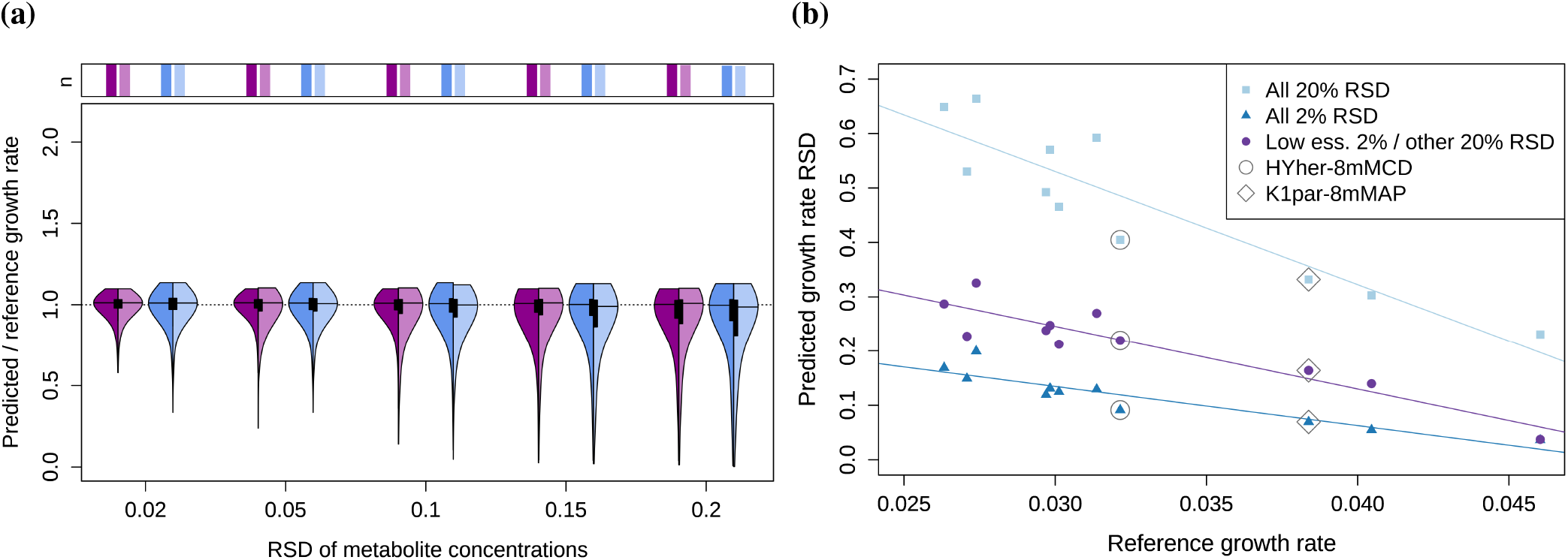
Panel a: the most important exchange rates for FBA predictions are cell line specific – the left halves show the predictions when the AAs with low normalized uptakes are chosen according to the cell line specific data; in the right halves they are chosen based on the data of the other cell line. In both cases, the RSDs of the chosen AAs are set to 2%, the remaining RSDs are varied. Purple: `K1par-8mMAP`, blue: `HYher-8mMCD`. For simplicity, the data is shown only for the sampling frequency of every 6 h. The barplots above the plot indicate the fraction of feasible FBA solutions. The apparent cutoffs on the top are artifacts due to the visualization. Panel b: Accurate measurements of cell line/condition specific low uptake AAs improve predictions in all datasets; the overall prediction accuracy gets worse at lower reference growth rates. The RSDs of metabolite concentrations were all set to 20% (“All 20% RSD”), 2% (“All 2% RSD”) or the low uptake AAs were set to 2% and the rest to 20% (“Low ess. 2% / other 20% RSD”). The lines represent linear fits.

Due to the fact that the most important AAs are cell line specific, we extended the analysis to eleven datasets from Széliová et al. ^20^ which include nine different cell lines, some of them in various media compositions (CD-CHO or ActiPro, 8 mM or 0 mM glutamine). We calculated normalized uptake rates for all datasets (for each specific biomass composition) and compared the lists of AAs with uptakes smaller than 1.5× of the biomass requirements. For all cell lines/conditions the list included lysine, phenylalanine and arginine, except for DXB11 datasets, where arginine was predicted to be nonessential (this is because no cell line specific model for DXB11 was available^9^ and arginine is not essential in the generic model). Isoleucine was not part of the low essential group in any of the datasets. For the remaining essential AAs, the normalized uptake rates varied among cell lines and conditions, but no consistent pattern was observed.

We ran FBAs for all the datasets, where we varied the RSD of all metabolites (2-20% RSD) or kept the RSDs low uptake essential AAs at 2% (and varied the rest). Figure 5 b shows that in all cases measuring the low uptake AAs accurately improves predictions. No cell line or condition specific effect was observed. However, we found a dependence on growth rate – the lower the reference growth rate, the worse the quality of the predicted growth rates. This is likely due to the fact that slower cells also have lower uptake rates, which in turn have bigger relative errors (as shown in the previous section).

### 3.4 | Accurate concentrations and high sampling frequency are needed for growth comparisons

Often, the goal of FBA is to compare (predicted) growth rates between different cell lines or conditions^32,17,33^. Therefore, we wanted to determine how often we are able to correctly identify the growth rate difference between two selected cell lines at the different sampling frequencies or RSDs of the concentration measurements. Again the difference from the reference FBAs was regarded as a “true” difference in the growth rates (0.0384 h^−1^ for `K1par-8mMAP` vs. 0.0321 h^−1^ for `HYher-8mMCD`, resulting in a 0.0063 h^−1^ or 16% difference). As described earlier (Section 3.2), we ran 1000 FBAs for the two cell lines at fixed sampling frequencies and metabolite concentration RSDs (see Figure 3). We compared the predicted growth rates of the two cell lines for all possible pairs (10^6^ pairs for each sampling frequency and each RSD). In some cases the FBAs for one or both cell lines had no feasible solution, so the total number of comparisons was lower (see legend in Figure A9 for the number of comparisons). In Figure A9 we normalized the predicted differences by the expected difference in growth rate (0.0063 h^−1^) and plotted them as cumulative distributions. The value of 1 represents the expected difference in growth rates. All values below zero mean that the predicted growth rate difference was opposite to what was expected – the faster-growing cell line had a smaller predicted growth rate than the slower-growing cell line.

As expected, with increasing RSDs of the concentration data and decreasing sampling frequency, there is a growing number of solutions that incorrectly identify the faster-growing cell line and the distributions get broader. For example at a daily sampling frequency and 20% RSD (Figure A9, right plot, green line), there are around 20% cases where the predicted difference is 5× higher than the “true” difference and almost 50% cases where the predicted difference is in the opposite direction.

Figure 6 a shows the (cumulative) probability of correctly predicting the faster-growing cell line. It can be used to determine the measurement accuracy and sampling frequency that are needed to identify the faster cell line in at least 80% of cases. If the concentration measurements are accurate enough, sampling frequencies of 6 h and 12 h lead to correct identification in at least 80% of cases. This probability quickly decreases with larger RSDs in the concentration measurements. In the worst case, less than 60% FBAs correctly predict which cell line grows faster, which is only slightly better than chance.

**FIGURE 6.**
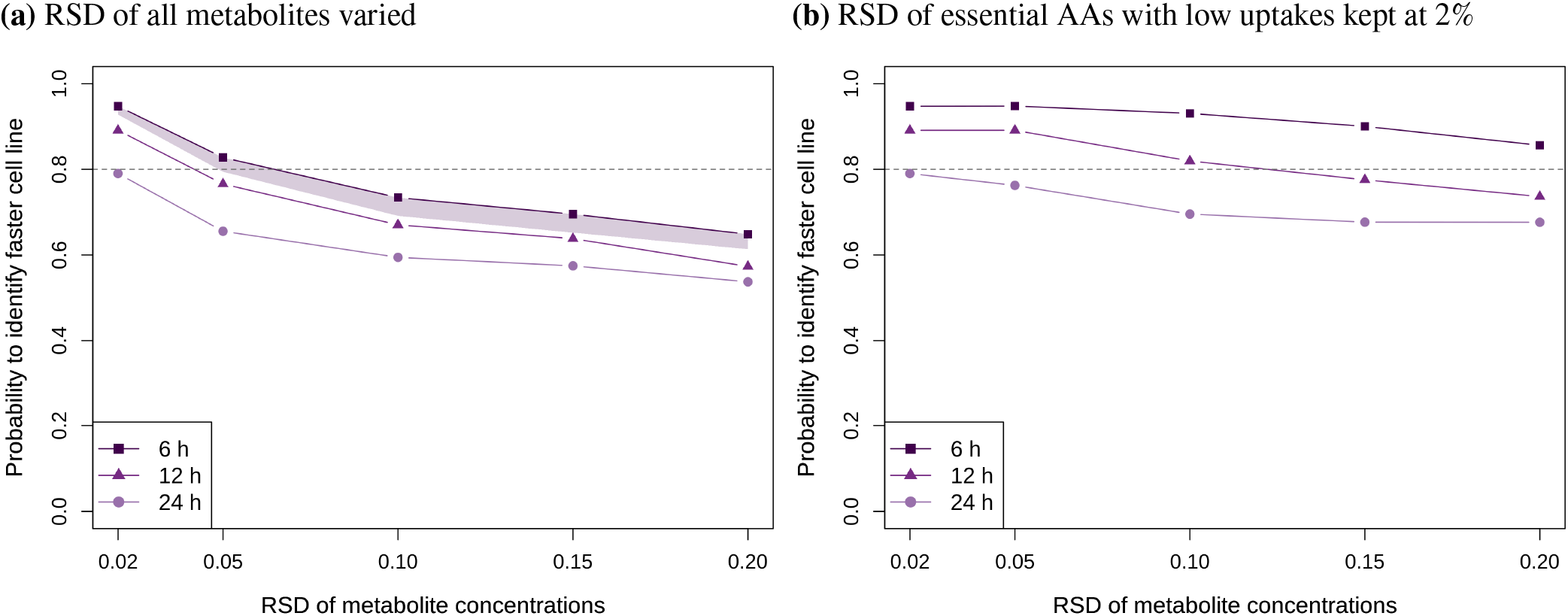
Percentage of FBAs where faster-growing cell line is correctly identified at different concentration RSDs and sampling frequencies. In panel a, concentration RSDs of all metabolites are varied between 2-20%. In panel b, the RSDs of low uptake essential AAs are kept at 2% (as in Figures 4 d and A6 d), while the RSDs of the remaining metabolites are varied between 2-20%. The shaded area in panel a represents results with different sampling schedules when 4 samples per day are taken: regular sampling every 6 h; every 4 h with a 12 h gap; every 2.5 h with a 16.5 h gap; every 12 h with 6 instead of 3 replicates. For the schedules with a gap, two versions were done with different positioning of the gaps, e.g. sampling at hours 0, 4, 8, 12, 24, 30, 36 etc. or 0, 12, 16, 20, 24, 36 etc.

We also checked whether it is necessary that the time points are equally spaced, as sampling every 6 h is impractical to do in a laboratory. Therefore, we simulated data with different sampling schedules where the sampling time points are close together during the (working) day (e.g. every 4 or 2.5 h), followed by a longer interval without sampling (12 or 16.5 h). In addition, we simulated sampling every 12 h, but with six rather then three replicates. The results for these alternative sampling schedules lie within the shaded area in Figure 6 a.

All previous simulations were done with three replicates at each time point (as illustrated in Figure 1). We wanted to test whether doubling the replicate number leads to a considerable improvement in the predictions of growth rate. Figure A10 a shows the probability of identifying the faster-growing cell line when the number of replicates is increased to six. The improvement compared to three replicates is very small. For comparison, we increased the number of replicates to 100 (Figure A10 b), although this is usually not feasible in a laboratory. The improvement is only modest, especially for the higher RSDs of metabolite-concentrations. If the RSD is 15% or bigger and sampling is done on a daily frequency, there is still a more than 20% chance to incorrectly identify the faster-growing cell line even with 100 replicates.

Finally, Figure 6 b shows the probability of correctly predicting the faster-growing cell line when the RSDs of essential AAs with low uptakes are kept at 2% and only the RSDs of the remaining AAs are varied (as in Figure 4 d and Figure A6 d, left halves). For the highest sampling frequency (every 6 h), the faster cell line is identified in at least 80% cases for all RSDs.

In the previous paragraphs we showed comparisons between the two selected datasets with 16% difference in growth rate. However, the size of the difference is another factor that affects the prediction accuracy. We expected that the bigger the growth rate difference, the easier it is to detect it. To verify this, we compared all possible pairs of the eleven datasets^20^, resulting in 55 comparisons, and plotted the probability to detect the faster cell line as a function of the growth rate difference. Two cases are shown as examples – 1. RSDs of all metabolite concentrations are 20% and 2. the low uptake AAs are measured accurately with 2% RSD and the rest with 20% RSD (Figure 7). Generally, the higher the growth rate difference, the easier it is to detect the faster cell line. However, for the case with 20% RSD of all the measurements, the difference cannot be reliably detected for most of the tested combinations, even at the biggest growth rate differences. Reducing the measurement error of the low uptake AAs leads to an improvement in the predictions and the faster cell line can be reliably detected if the difference is at least 20%. Small growth rate differences (0.4-10%) could not be reliably detected in any of the cases.

**Figure 7.**
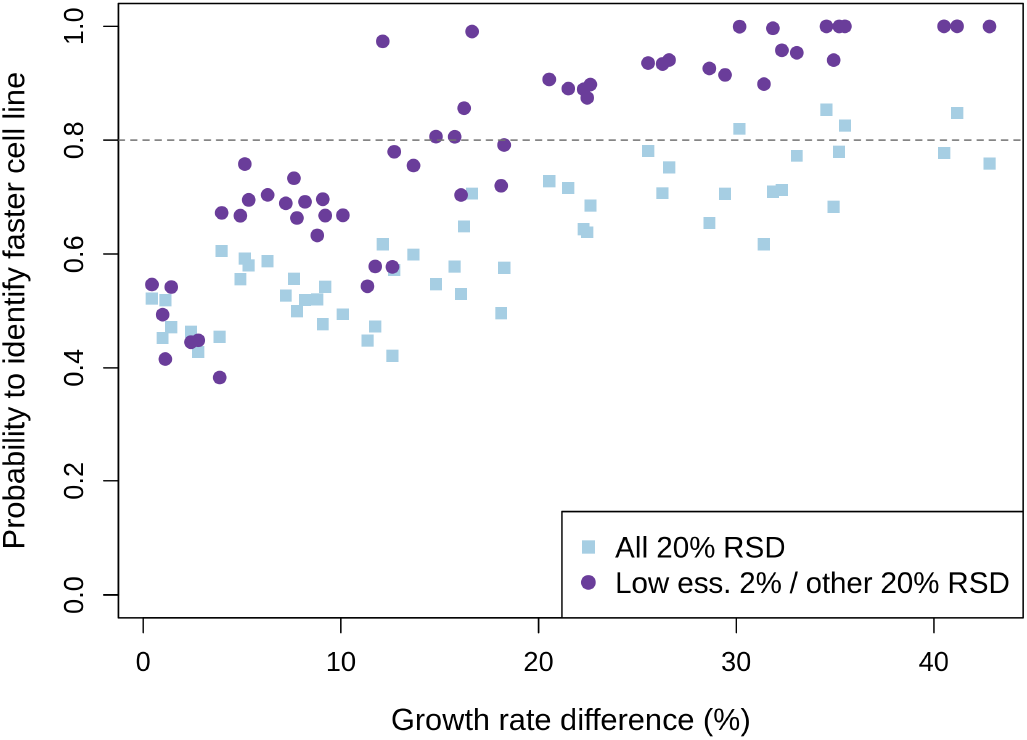
Probability to identify faster cell line as a function of growth rate difference at sampling frequency of every 6 h for two cases: “All 20% RSD” – RSDs of concentration measurements of all metabolites are 20%; “Low ess. 2% / other 20% RSD” – RSDs of low essential AA concentrations are 2%, the rest 20%.

## 4 | DISCUSSION

Systems level analysis of CHO cells is needed in order to design rational engineering and optimization strategies to streamline cell line and process development. The genome-scale metabolic model of CHO^9^ provides a basis for constraint-based metabolic modeling methods such as FBA. To successfully use these and obtain useful predictions, it is necessary to feed the model with accurate data, mainly uptake and secretion rates of extracellular metabolites. CHO cells are cultivated in complex media and consume or secrete numerous metabolites, including glucose, lactate, ammonium and all amino acids. Typically, more than 20 metabolite exchange rates have to be determined as an input for FBA.

Previously, we showed that accurate quantification of uptake and secretion rates is essential for good predictions of CHO growth rates by FBA; to achieve the required accuracy it is necessary to analyze a sufficient number of time points throughout the relevant culture phases^20^. However, we did not systematically analyse the effect of measurement error and sampling frequency on the rate calculations and growth rate predictions, nor did we identify which metabolites have the biggest impact. These points are addressed here.

In Figure 2 we showed examples of error propagation from concentration data to the calculated uptake rates. The smaller the uptake rate, the higher the relative error of the rate, which was already pointed out by Hädicke et al.^34^. The analysis also shows the importance of a sufficiently high sampling density throughout the culture. The comparisons in Figures 2 demonstrate that sampling only once per day (or 4 time points during the exponential phase, as is standard practice) leads to errors in the rates that are several times larger than the errors of the concentration data. There are also more extreme values (e.g. very high/low rates or even rates that point in the opposite direction) which are basically unusable for any meaningful analysis with FBA. This is nicely demonstrated in Figure 3 by the big variance and the high number of infeasible solutions. Consequently, insufficient measurement accuracy as well as low sampling frequency also make it almost impossible to reliably compare the predicted growth rates of two cell lines (Figure 6).

There are more than 20 uptake and secretion rates that are used as inputs for FBA. However, their impact on the predictions of growth rates are vastly different. We expected that the prediction accuracy might improve if we measure those metabolites accurately that have the biggest relative errors of the rates. Even though this approach led to a big improvement in the predictions, further improvement was observed when we considered the biological roles of the metabolites. Even though all AAs are used as building blocks for the biomass, there are big differences in their exchange rates and metabolism. Some AAs can be synthesized in CHO, while others are essential and need to be taken up from the medium. Most of the essential AAs are consumed at very low rates and are predominantly used for the synthesis of the biomass but some can also be partially catabolized. We expected that the amino acids which are solely used for the biomass synthesis would have the biggest impact on the growth rate predictions, which was confirmed by our analysis in Figure 4. Interestingly, the uptake rates of glucose and glutamine, which are the main energy sources for CHO^35^, have no impact on the growth rate predictions, suggesting that energy provision is not the limiting factor for the predicted growth rates. However, this only applies to the cultures supplemented with an excess of energy sources, which was the case for all the batch cultivations in Széliová et al.^20^. If energy sources in the medium were insufficient, their uptake rates would likely become more important for predictions of growth rate.

A similar analysis was done by Goudar et al. ^36^ where error propagation from metabolite concentrations to rates and then to metabolic fluxes was analysed and revealed that lesser metabolic fluxes (AA metabolism) were strongly influenced by the errors in the greater exchange rates (e.g. glucose, lactate), but not by the lesser exchange rates (AAs). In contrast, our analysis showed that low exchange rates of essential AAs had the biggest impact on the predictions of growth rate (which was not analysed in Goudar et al.^36^). Among other differences between the studies is that in Goudar et al. a small metabolic network was used and not all relevant nutrient exchange rates were considered (e.g. all AAs). However, in genome-scale models, more than 20 exchange rates are typically used, which leads to a bigger complexity of the analysis. Furthermore, the experiments were done in perfusion cultures, where the procedure for determination of rates is different from batch cultivation, since they are in steady state. Nevertheless, both of these studies complement each other by showing the importance of the accurate rate determination for metabolic modelling.

An important point brought up in a study by Bayer et al.^37^ was that the choice of the rate calculation method largely impacts the accuracy of the calculated rates. They concluded that fitting a cubic smoothing spline function is much better than doing a stepwise integration which is more sensitive to experimental errors. Because in this work we analysed data from the exponential phase of batch cultivations, an exponential function was deemed more appropriate than a spline function. Similar to the spline function, an exponential function is fitted to the data from the whole process and therefore should not be influenced by experimental noise as much as step-wise integration.

In conclusion, our analyses show that to obtain accurate FBA predictions of growth rates, the essential amino acids with low uptake rates should be quantified with the highest possible accuracy and at a sufficient sampling frequency throughout the culture. These aspects should be taken into consideration when planning experiments. The RSDs simulated in this paper represent the overall variation among replicates. This can be divided into biological variation and the variation stemming from the measurement errors. Hädicke et al. ^34^ provided a guide for how the contributions of these errors to the total variance can be analysed. Generally, the biological variance is bigger than the measurement variance and cannot be influenced, but it can be estimated from historical data, if available. On the other hand, measurement variance can be reduced by choosing an appropriate method that can accurately quantify the AAs that have the biggest impact on predictions and considering all other aspects that could impact the final data quality (e.g. storage of the samples at −80 °C immediately after sampling, avoiding freeze-thawing etc.). The measurement accuracy of six example amino acids (important for `K1par-8mMAP`) is discussed in the supplementary section A1.

When planning experiments, simulations similar to the ones presented in this paper can be performed to determine whether a certain accuracy and sampling frequency are sufficient to answer the required scientific questions. Even though the growth rate difference between two conditions/cell lines is not known beforehand, researchers can define what difference they want to be able to detect and adjust the experimental design accordingly. Realistically, according to Figure 7, growth rate differences below 15% are unlikely to be detected by FBA even if high effort goes into performing an experiment.

This work showed the effect of measurement error and sampling frequency on the predictions of growth rate and highlights the difficulty of obtaining accurate uptake rates of the AAs with low uptakes even when the experimental errors are low. However, these results are specific to the use of biomass objective function, which is one of the most common objective functions^38^. In a recent preprint, Schinn et al. ^39^ report that the objective function has a big impact on the prediction accuracy and the biomass objective function was correlated with poor predictions. A suggested alternative was minimization of cytosolic NADPH regeneration, which was correlated with good predictions. In another work, Chen et al. ^40^ proposed using minimization of the uptake of a nonessential metabolite (e.g. glucose) as the objective function. In their approach, the growth rate is not predicted, but fixed to the experimental value and the uptake rates of essential AAs are predicted (based on the biomass requirements). As a result, the experimental error of these AAs would not affect the FBA solution.

## 5 | ACKNOWLEDGEMENTS

This work has been supported by the COMET center “acib: Next Generation Bioproduction”, which is funded by BMK, BMDW, SFG, Standortagentur Tirol, Government of Lower Austria and Vienna Business Agency in the framework of COMET - Competence Centres for Excellent Technologies. The COMET-Funding Program is managed by the Austrian Research Promotion Agency FFG. Additional funding came from the PhD program BioToP (Biomolecular Technology of Proteins) of the Austrian Science Fund (FWF Project W1224).

## Abbreviations

AAs: amino acids
CHO: chinese hamster ovary
FBA: flux balance analysis
mAb: monoclonal antibody
RSD: relative standard deviation
RSE: relative standard error
SE: standard error

