## Supplementary material for "Error propagation in constraint-based modeling of Chinese hamster ovary cells"

**How to cite this article:** Diana Széliová, Dmytro Iurashev, David E Ruckerbauer, Gunda Koellensperger Nicole Borth, Michael Melcher, and Jürgen Zanghellini (2020), Error propagation in constraint-based modeling of Chinese hamster ovary cells, *Biotechnology Journal*, *2020;00:1–6*.

### APPENDIX

**TABLE A1** Initial concentration $[\hat{i}]_0$ of the medium components (mM) / biomass $\widehat{\mathrm{BM}}_0$ (g L$^{-1}$) and exchange rates $\hat{q}_i$ (mmol g$^{-1}$ h$^{-1}$) for two selected cell lines `K1par-8mMAP` (top half) and `HYher-8mMCD` (bottom half). For biomass, the column $\hat{q}_i$ corresponds to growth rate $\hat{\mu}$ (h$^{-1}$) of the reference FBA.

| $i$ | $[\hat{i}]_0$ | $\hat{q}_i$ | $i$ | $[\hat{i}]_0$ | $\hat{q}_i$ | $i$ | $[\hat{i}]_0$ | $\hat{q}_i$ | $i$ | $[\hat{i}]_0$ | $\hat{q}_i$ | $i$ | $[\hat{i}]_0$ | $\hat{q}_i$ |
|---|---|---|---|---|---|---|---|---|---|---|---|---|---|---|
| Ala | 0.165 | 0.07 | Glc | 32.262 | -0.523 | Ile | 3.292 | -0.016 | $NH_4$ | 3.206 | 0.167 | Trp* | 1.019 | -0.002 |
| Arg | 1.488 | -0.013 | Gln | 8.591 | -0.225 | Lac | 4.313 | 0.756 | Phe | 1.571 | -0.008 | Tyr | 0.891 | -0.007 |
| Asn | 5.314 | -0.07 | Glu | 2.668 | 0.03 | Leu | 5.164 | -0.026 | Pro | 6.632 | -0.011 | Val | 3.926 | -0.016 |
| Asp | 1.819 | 0.041 | Gly | 1.384 | 0.029 | Lys | 2.707 | -0.017 | Ser | 6.236 | -0.056 | | | |
| Cys* | 0.379 | -0.004 | His | 1.134 | -0.006 | Met | 1.366 | -0.006 | Thr | 3.646 | -0.01 | BM | 0.040 | 0.038 |
| Ala | 0.151 | 0.036 | Glc | 34.831 | -0.441 | Ile | 2.752 | -0.011 | NH4 | 0.797 | 0.114 | Trp* | 1.019 | -0.001 |
| Arg | 2.081 | -0.012 | Gln | 8.906 | -0.101 | Lac | 7.047 | 0.477 | Phe | 1.330 | -0.008 | Tyr | 0.985 | -0.007 |
| Asn | 5.999 | -0.087 | Glu | 1.826 | 0.013 | Leu | 4.048 | -0.020 | Pro | 4.349 | -0.020 | Val | 3.150 | -0.017 |
| Asp | 1.381 | 0.020 | Gly | 0.128 | 0.019 | Lys | 3.016 | -0.016 | Ser | 5.138 | -0.046 | | | |
| Cys* | 0.379 | -0.004 | His | 1.198 | -0.006 | Met | 0.875 | -0.004 | Thr | 3.016 | -0.013 | BM | 0.045 | 0.032 |

Source: Széliová et al.[20]. The full dataset for all cell lines/conditions is available at Data Mendeley (https://data.mendeley.com/datasets/5vn5m33wpr/draft?a=3e7afba4-42e0-45b4-9e16-7adb419251f9).
* rates of Cys and Trp were predicted from the reference FBA. The initial concentrations are from the CD-CHO medium patent.

### A1 REMARKS ON QUANTIFICATION OF AMINO ACIDS BY MASS SPECTROMETRY

Accurate absolute quantification of amino acids by mass spectrometry (MS) involves isotope dilution, regardless whether GC-MS (gas chromatography-mass spectrometry) or LC-MS (liquid chromatography–mass spectrometry) methods are implemented. Internal standardization is accomplished either by $^{13}$C enriched or deuterated standards. Isotopic labelling of biomass proved to offer cost-effective pipelines for standard production[41]. In fermentation media, protein precipitation and efficient amino acid extraction is ensured by using polar solvents, both a prerequisite for accurate quantification. The internal standard is added as early as possible upon extraction, in order to compensate for losses during sample preparation (to ensure ideally 100% recovery). In this way, experimental precision in the low percent range can be achieved on a routine basis.

The accuracy of amino acid analysis in fermentation media can be validated by using the certified reference material SRM 1950, human plasma, as the same protocols for sample preparation and MS methods can be applied. While threonine and lysine can be accurately quantified by both platforms, LC-MS and GC-MS, respectively, accurate quantification of arginine, cysteine, phenylalanine and tryptophan demands LC-MS based analysis.

Using orthogonal GC-MS and LC-MS, threonine and lysine concentrations could be certified in the certified reference material for human plasma, SRM 1950, with expanded uncertainties of 5 and 10% respectively. The expanded uncertainty covers the measurand with approximately 95% confidence. Thus it expresses both the observed difference between the results from the LC-MS and GC-MS methods and their respective uncertainties, in addition to uncertainty components related to purity of the standards used for calibration. Using LC-MS methods alone, excellent experimental precisions ranging at 2-3% are technically feasible. The repeatabilities given in table A2 were observed by analysing independently prepared plasma samples[42] by hydrophilic interaction chromatography (based on a zwitterionic stationary phase at pH 9) providing isomer separation of threonine and homoserine.

Decomposition upon derivatization in GC-MS hampers the analysis of phenylalanine, arginine and cysteine. Despite internal standardization and robotic just-in-time derivatization, otherwise ensuring accuracy and precision, the detrimental decomposition processes result in poor recoveries and high uncertainties for these amino acids. Accordingly, for these amino acids only reference values based on LC-MS analysis were reported in SRM 1950. The expanded uncertainties ranged between 3 and 15%. Phenylalanine revealed an expanded uncertainty of 15% based on 3 independent LC-MS protocols. Using only one method as in the case of arginine, an expanded uncertainty of 3% was obtained, which corresponds to typical LC-MS precisions observed in other laboratories[42] (see table A2). The poor precision as obtained for cysteine in SRM 1950 can be improved, as in the certification campaign, no derivatization was implemented. In fact, accurate analysis of cysteine, containing a highly reactive thiol group, requires chemical protection during extraction. A protocol using N-ethylmaleimid protection[43] of the reactive thiol improved the precision achievable by LC-MS (5-10%).

Tryptophan is known to be a highly unstable compound, making proper standardization a challenge as such. This is exemplified through the fact that neither the certified reference standard for amino acids (NIST SRM 2389a) nor the human plasma SRM 1950 (see Table A2) gives concentrations for tryptophan. Special care has to be taken for the preparation of tryptophan standards. In GC-MS, accuracy is additionally jeopardized by the derivatization step. For calibration, tryptophan standards have to be prepared separately to the otherwise used SRM 2389a and freshly. Time between sampling, sample preparation and measurement has to be minimized. Taking all these measures, experimental repeatabilities of around 3% are achievable when using HILIC-MS and a minimum sample preparation (without preconcentration step by vacuum evaporation) [Mate Rusz, unpublished data].

**TABLE A2** Uncertainties of six amino acids important for cell line `K1par-8mMAP`

| Amino acid | Standard uncertainty % LC-MS (HILIC/RP) standards: SRM 2389a, tryptophan, N=5; 24 h isotope dilution using $^{13}$C yeast as standard | Standard uncertainty % LC-MS (HILIC), plasma, protein precipitation, $\mu$M concentrations, N=5 independently prepared samples isotope dilution using $^{13}$C yeast as standard | Expanded uncertainty % Certificate SRM 1950, NIST Human plasma reference material* isotope dilution using $^{13}$C yeast as standard |
|---|---|---|---|
| Arginine | 2 | 3 | 3 |
| Cysteine** | 4 | 5-10 | 15 |
| Lysine | 2 | 3 | 10 |
| Phenylalanine | 1-2 | 4 | 13 |
| Threonine*** | 2-3 | 3 | 5 |
| Tryptophan | 2 | 3 | No certified/no indicative value given |

* SRM 1950: for lysine and threonine, certified concentrations are given in SRM 1950 (orthogonal GC-MS, LC-MS measurements available using isotope dilution); for arginine, cysteine and phenylalanine only reference values are given since only LC-MS data (using isotope dilution) were available; no data for tryptophan. The uncertainty provided with each value is an expanded uncertainty about the mean to cover the measurand with approximately 95% confidence; it expresses both the observed difference between the results from the methods and their respective uncertainties, in addition to Type B components related to purity of the standards used, consistent with the ISO/JCGM Guide.

** accurate quantification of cysteine requires derivatization upon extraction. NEM derivatization can be accomplished during the extraction/protein precipitation step[43].

*** The isomers threonine/homoserine are not always separated, only by zicHILIC.

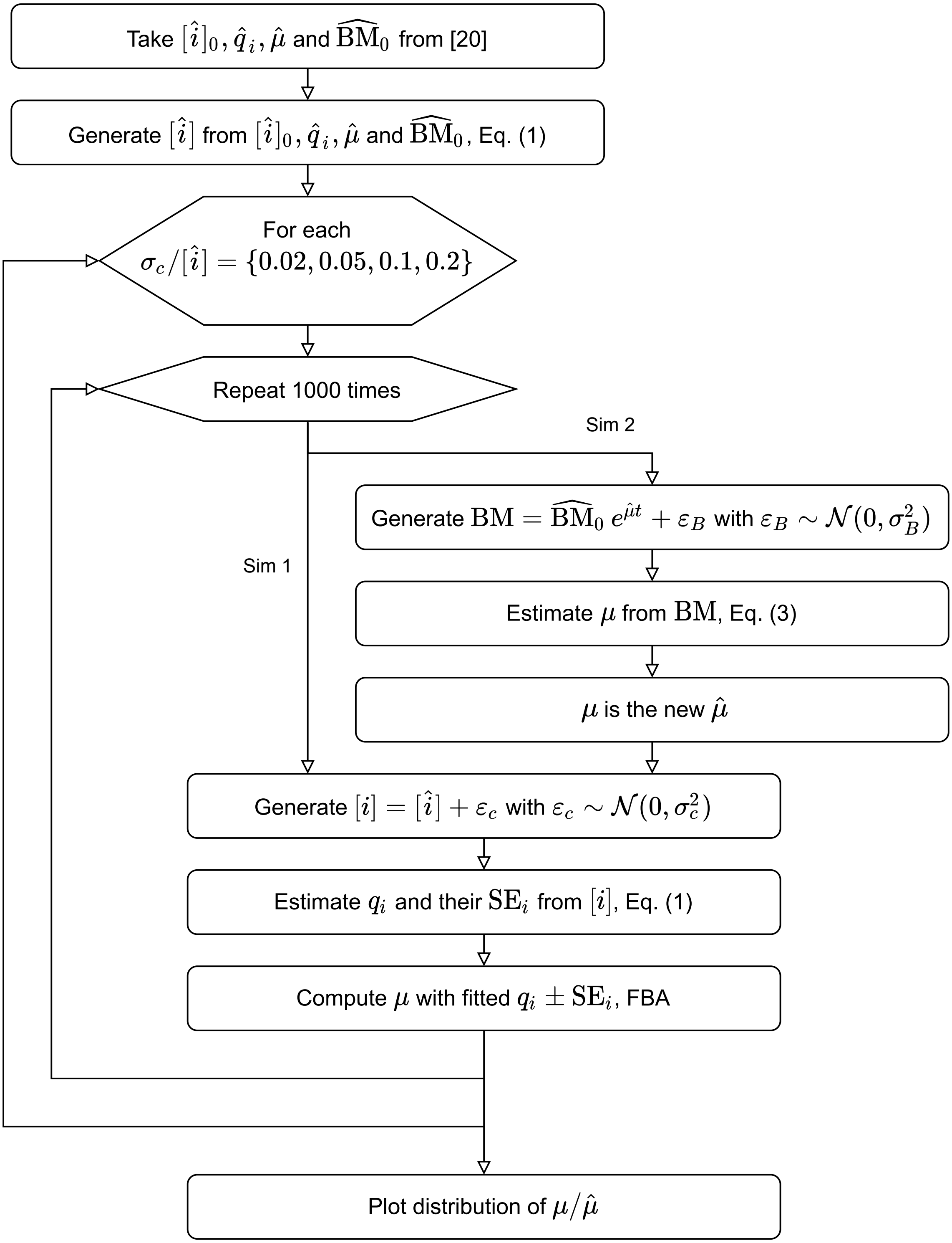


**FIGURE A1** Workflow of the simulations.

**(a)** K1par-8mMAP **(b)** HYher-8mMCD

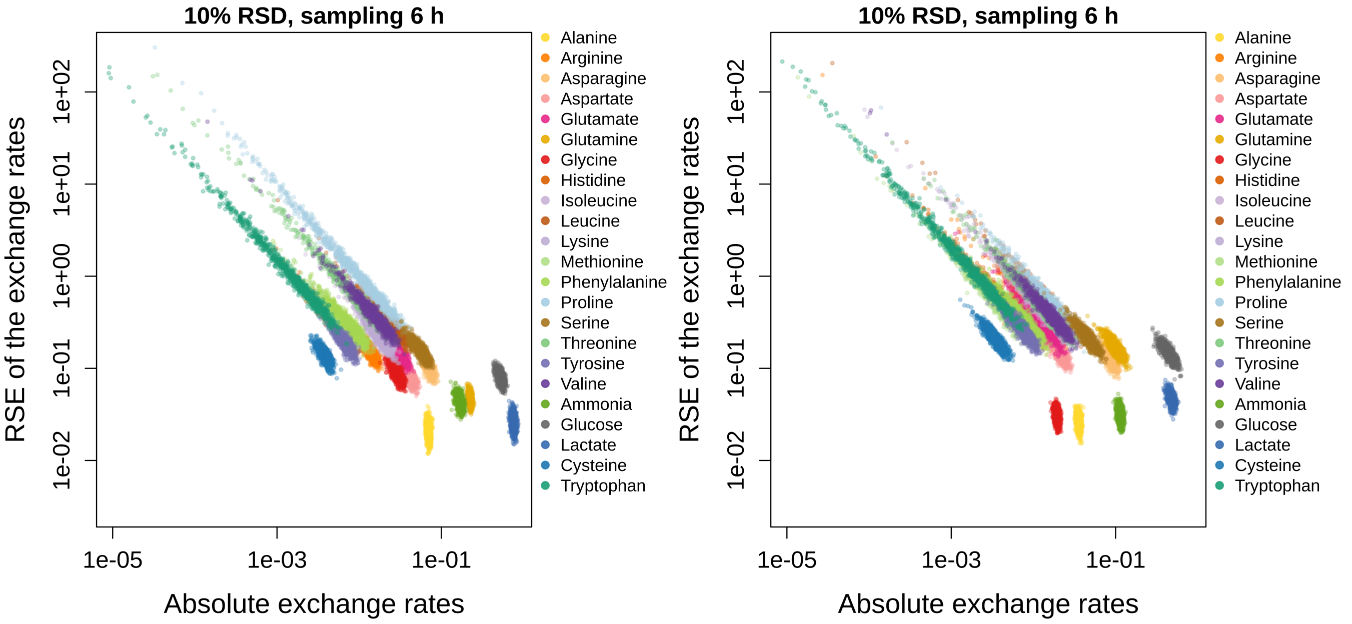


**FIGURE A2** Related to Figure 2 . RSEs of the exchange rates as a function of the absolute values of exchange rates for 10% RSD of the concentration data for cell lines `K1par-8mMAP` (panel a) and `HYher-8mMCD` (panel b).

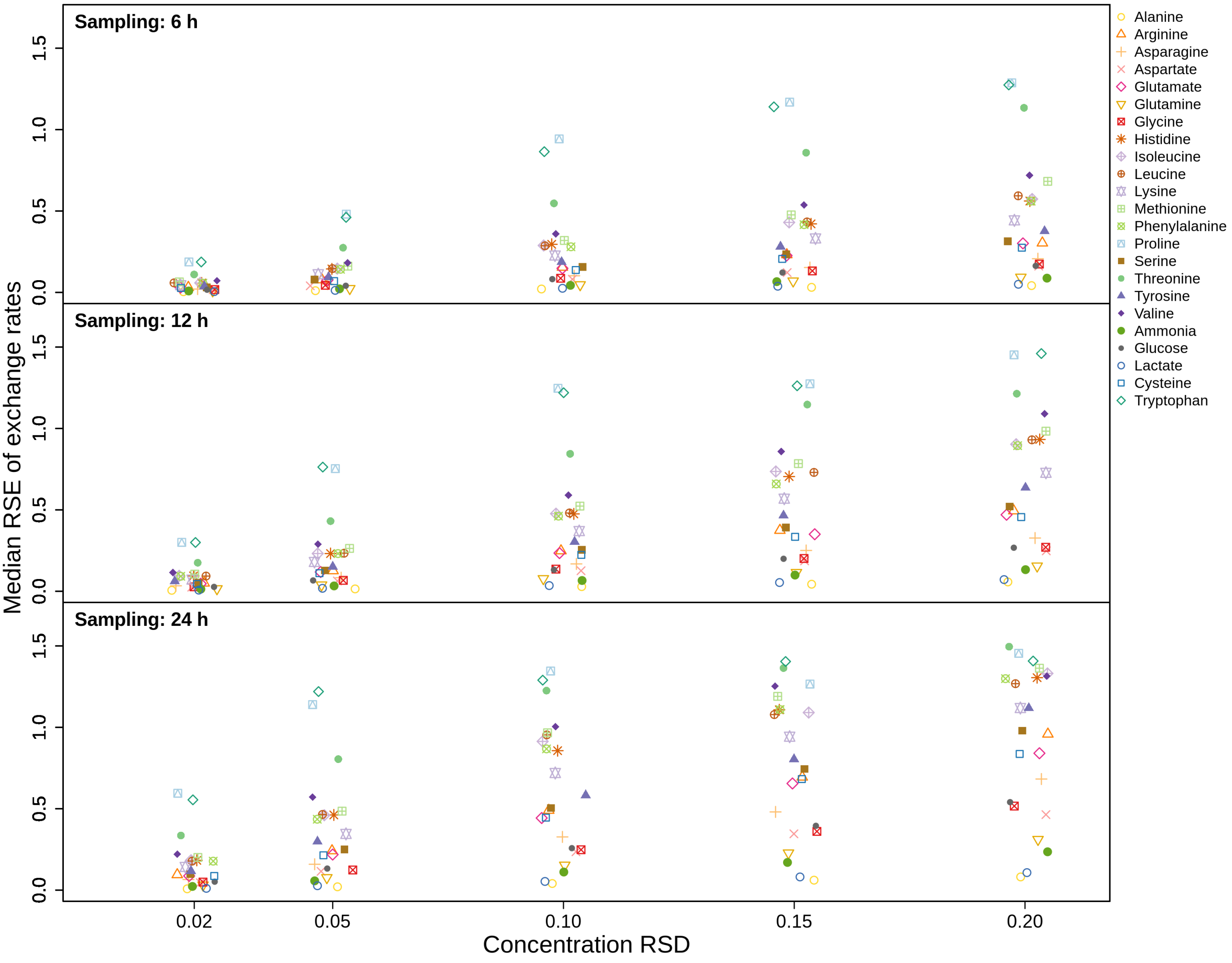


**FIGURE A3** Relationship of the concentration RSDs and median RSEs of the exchange rates for cell line `K1par-8mMAP`.

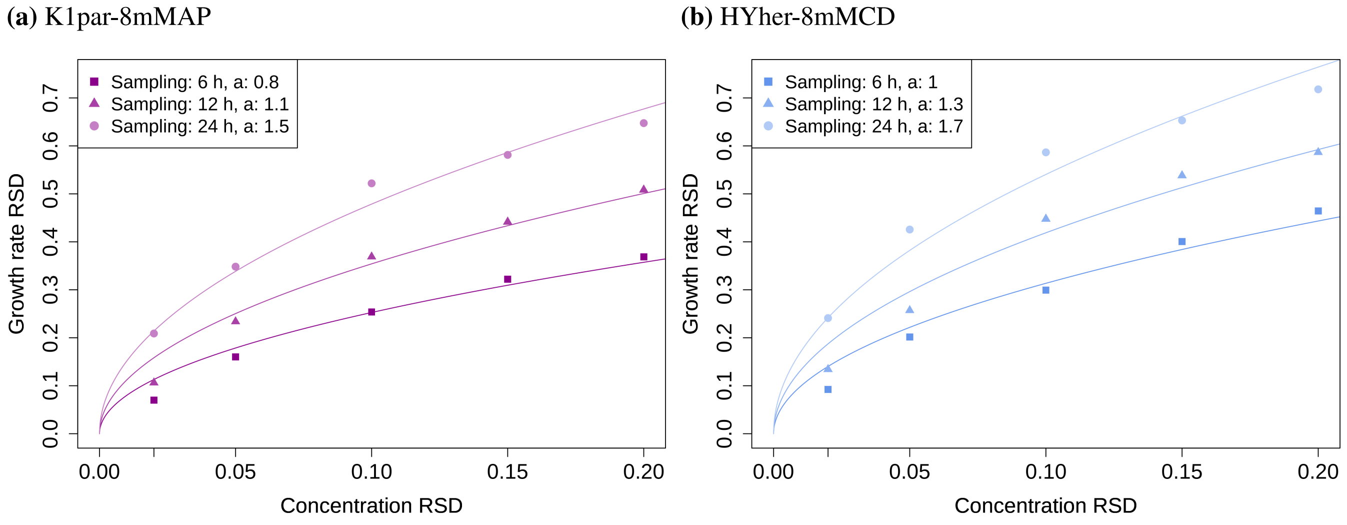


**FIGURE A4** Related to Figure 3 . RSDs of growth rate predictions as a function of concentration RSDs at different sampling frequencies. The growth rate RSDs were fitted with the function $a\sqrt{x}$, where $x$ are the concentration RSDs and $a$ is a fitting parameter (the fitted values are shown in the legend). Panel a shows results for the cell line `K1par-8mMAP`, panel b for `HYher-8mMCD`.

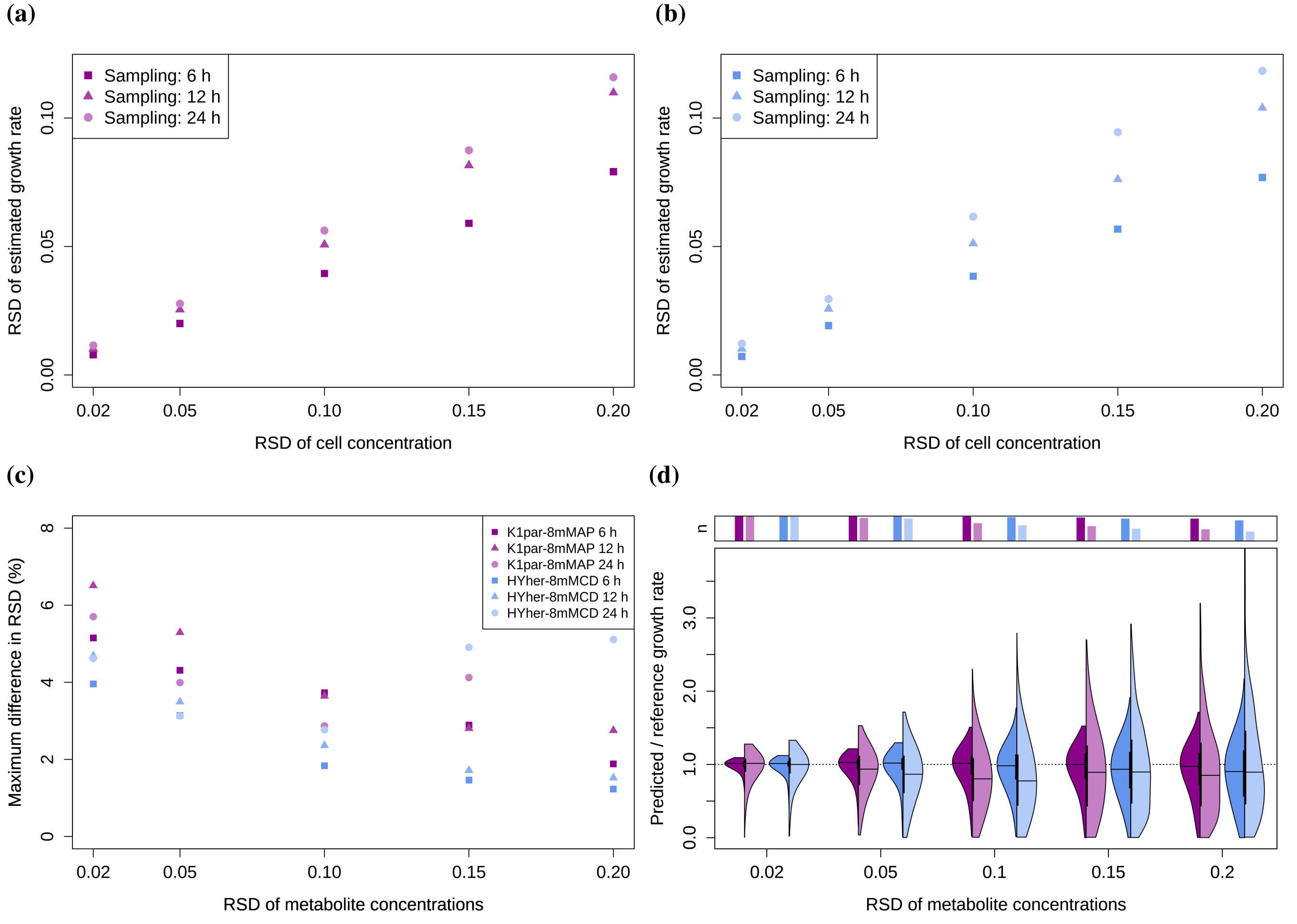


**FIGURE A5** Effect of the cell concentration RSD on the growth rate estimated with Equation (3) for `K1par-8mMAP` (panel a) and `HYher-8mMCD` (panel b). Panel c shows the maximum observed difference between the exchange rates estimated with Equation (1) with constant growth rate or growth rate estimated from cell concentrations perturbed with 6% RSD. Panel d: FBA predictions at different concentration RSDs for two cell lines (purple: `K1par-8mMAP`, blue: `HYher-8mMCD`). Compared to Figure 3 , not only metabolite concentrations, but also the cell concentrations are perturbed (with RSD of 6%). The left halves of the violin plots correspond to sampling every 6 h, the right halves every 24 h. The barplots above the plots indicate the fraction of feasible FBA solutions. The apparent cutoffs on the top at lower RSDs are artifacts due to the visualization.

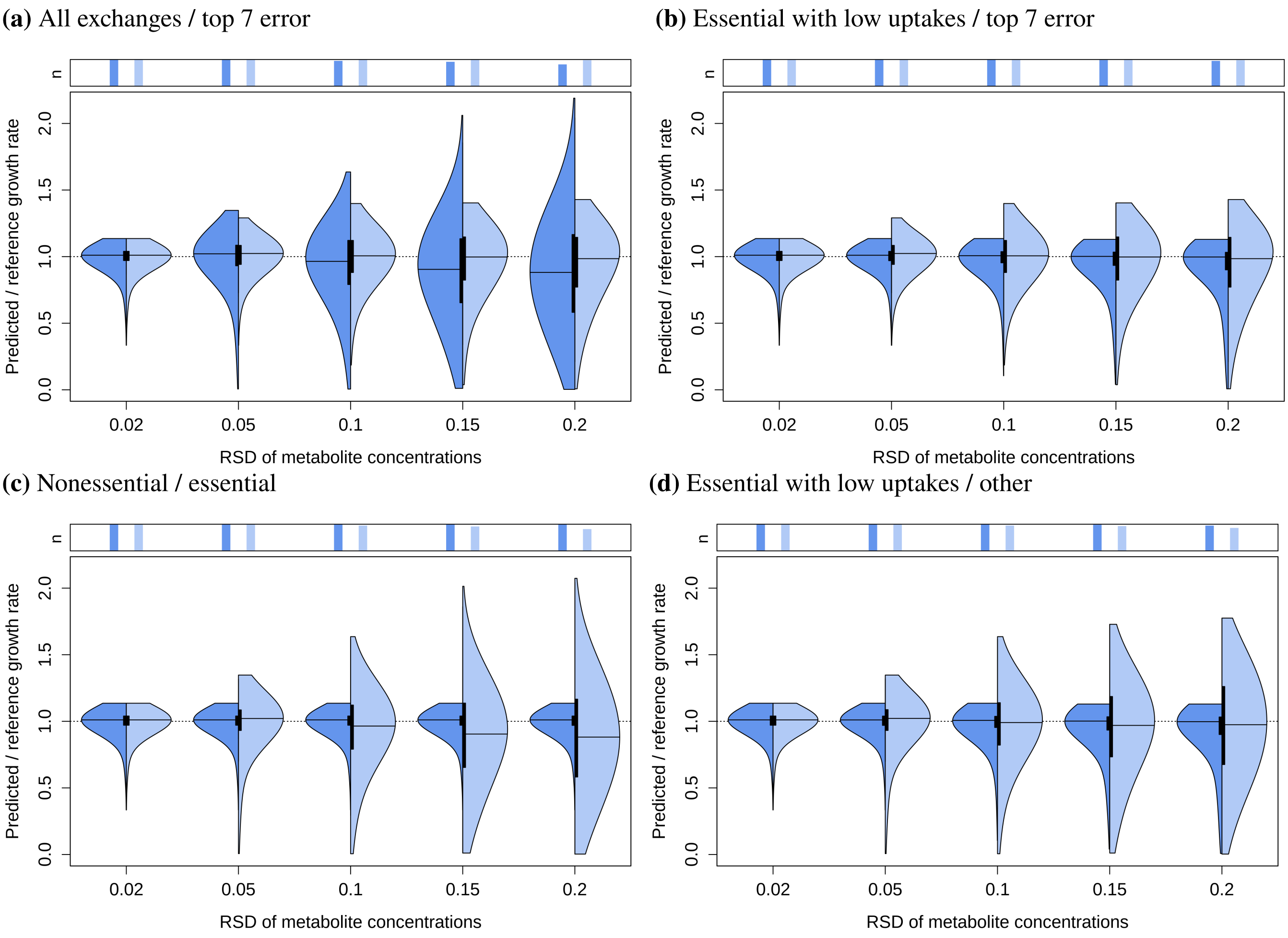


**FIGURE A6** Related to Figure 4 . FBA predictions when different groups of metabolites are measured accurately. In both panel a and b, the right halves of the violin plots show results when the RSDs of the metabolites with top 7 highest RSEs of the rates are set to 2% (tryptophan, isoleucine, threonine, methionine, proline, leucine, histidine). In panel a, the left halves show results when RSDs of all metabolites are varied (the same as the left halves in Figure 3 ) and in panel b when RSDs of the AAs with low normalized uptakes are set to 2%. In panel c, left halves, the RSDs of the essential AA concentrations are set to 2%, while all others are varied (group 1 constant/group 2 varied); in the right halves, the other way around (group 1 varied/group 2 constant). In panel d, left halves, the RSDs of essential AAs with uptakes $< 1.5\times$ biomass requirements (arginine, leucine, lysine, methionine, phenylalanine, tryptophan, cysteine) are set to 2%, while all others are varied (group 1 constant/group 2 varied; the same as in panel b left halves); in the right halves, the other way around (group 1 varied/group 2 constant). For simplicity, the data is shown only for cell line `HYher-8mMCD` at the sampling frequency of every 6 h. The barplots above the plots indicate the fraction of feasible FBA solutions. The apparent cutoffs on the top are artifacts due to the visualization.

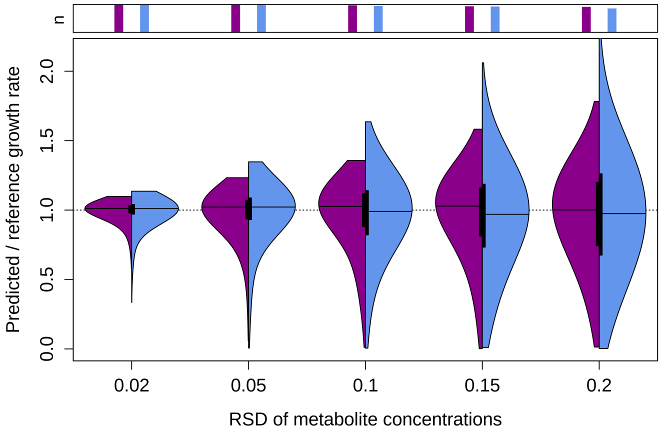


**FIGURE A7** Related to Figure 4 d and Figure A6 d. FBA predictions when the RSDs of essential AAs with low uptakes (< 1.5× biomass requirements) are varied (same as in the right halves of Figure 4 d and Figure A6 d) and all other uptakes are left unconstrained. Purple: `K1par-8mMAP`, blue: `HYher-8mMCD`. For simplicity, the data is shown only for the sampling frequency of every 6 h. The barplots above the plot indicate the fraction of feasible FBA solutions.

**(a)** Grouping 1, group 1 varied

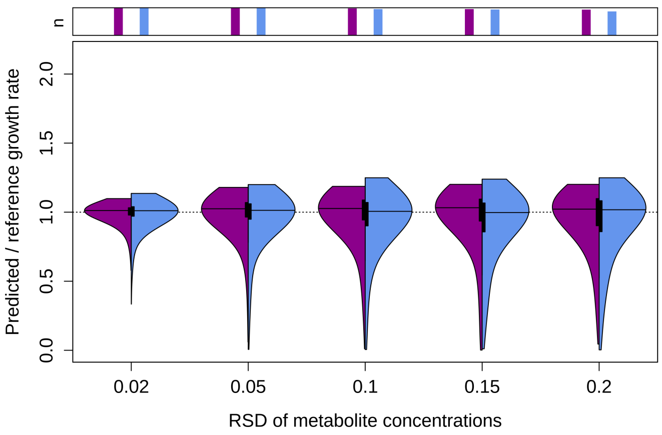


**(b)** Grouping 1, group 2 varied

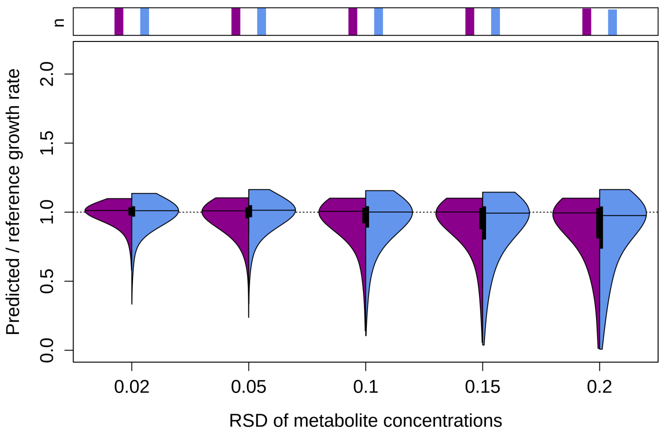


**(c)** Grouping 2, group 1 varied

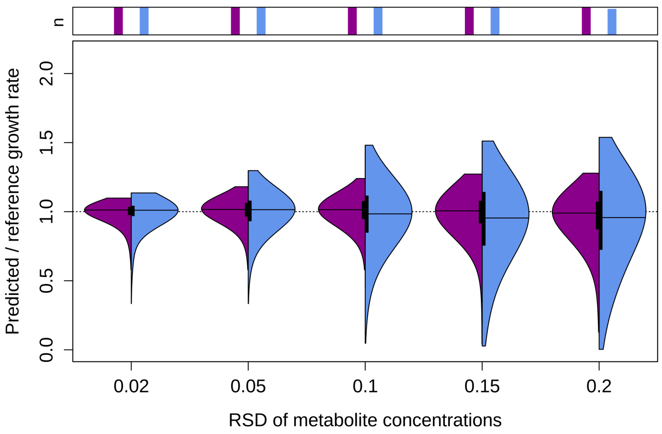


**(d)** Grouping 2, group 2 varied

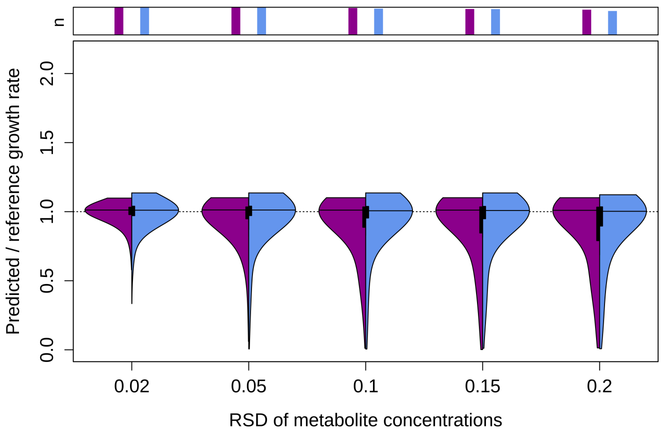


**(e)** Grouping 3, group 1 varied

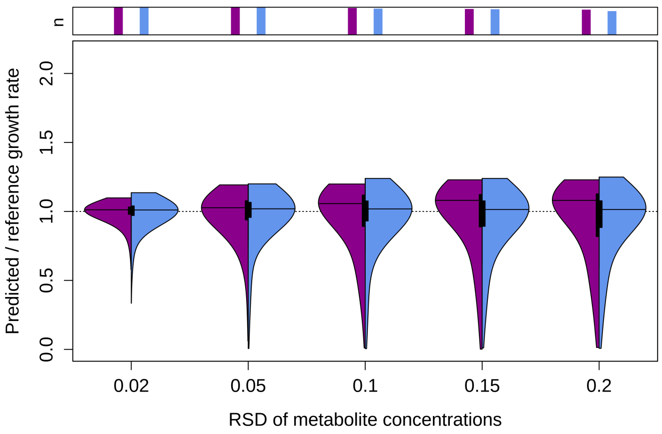


**(f)** Grouping 3, group 2 varied

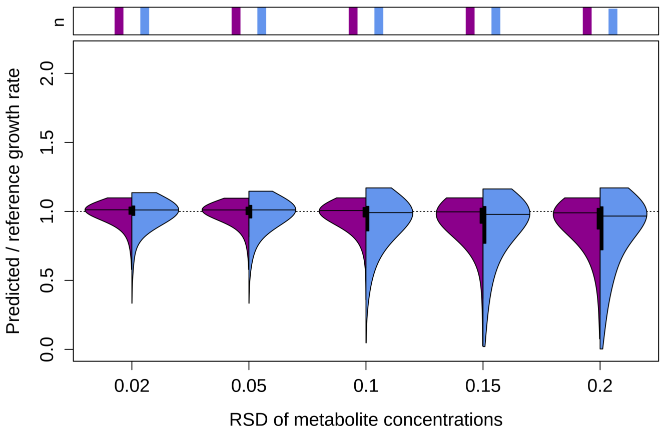


**FIGURE A8** Related to Figure 4 . The metabolites are randomly split into two groups and the RSDs of group 1 are varied, while keeping the RSDs of group 2 at 2% (and the other way around). The random grouping was repeated 3 times for each cell line (labelled as Grouping 1-3). Purple: `K1par-8mMAP`, blue: `HYher-8mMCD`. For simplicity, the data is shown only for the sampling frequency of every 6 h. The barplots above the plot indicate the fraction of feasible FBA solutions.

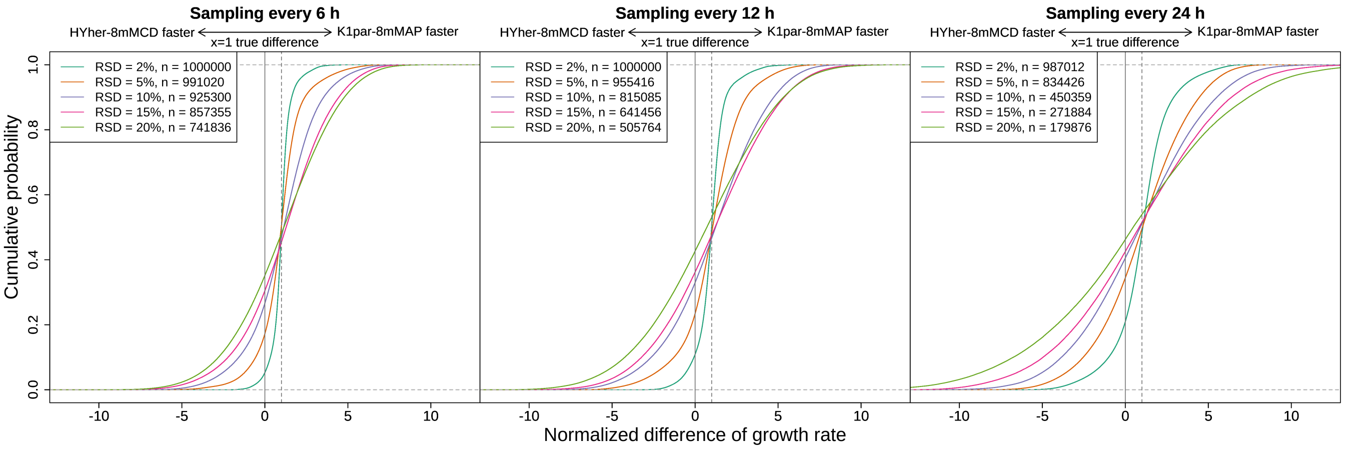


**FIGURE A9** Related to Figure 6 a. The differences in growth rates predicted by FBA for two cell lines were normalized by the expected difference and plotted as a cumulative distribution. "RSD" refers to the error of the metabolite concentration data and "n" indicates the number of comparisons where FBA solutions were feasible for both cell lines.

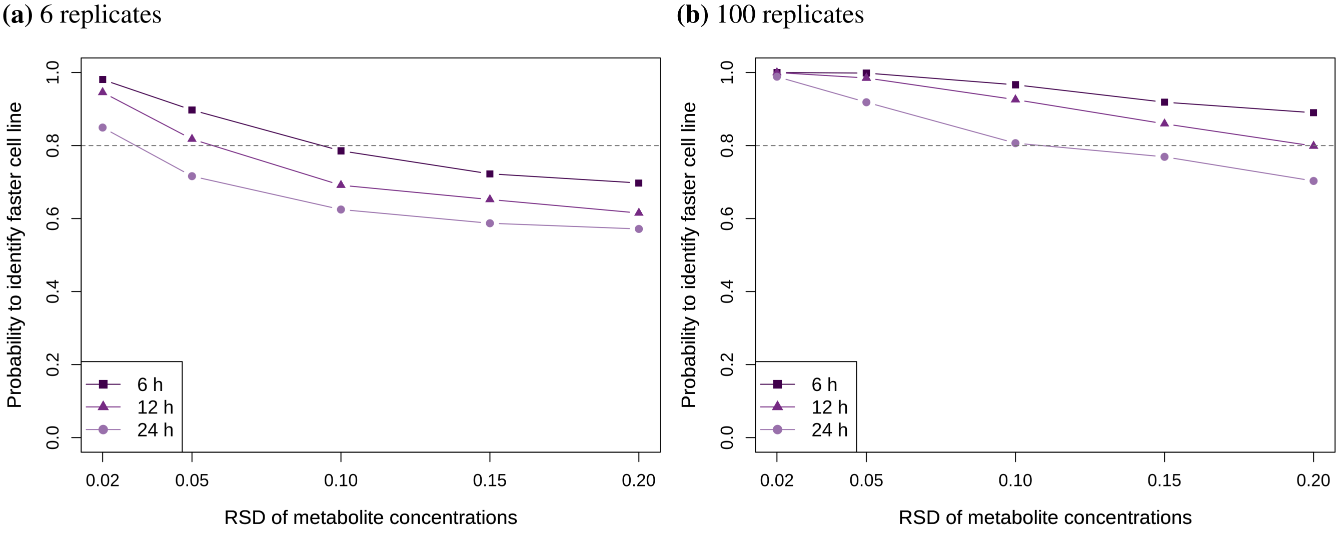


**FIGURE A10** Percentage of FBAs where faster growing cell line is correctly identified when 6 (panel a) or 100 (panel b) replicates are analysed at each time point (instead of 3 replicates as in Figure 6 a).
